# Cellular abundance shapes function in piRNA-guided genome defense

**DOI:** 10.1101/2021.03.04.433912

**Authors:** Pavol Genzor, Parthena Konstantinidou, Daniel Stoyko, Amirhossein Manzourolajdad, Celine Marlin Andrews, Alexandra R. Elchert, Constantinos Stathopoulos, Astrid D. Haase

## Abstract

Defense against genome invaders universally relies on RNA-guided immunity. Prokaryotic CRISPR/Cas and eukaryotic RNA interference pathways recognize targets by complementary base-pairing, which places the sequences of their guide RNAs at the center of self/nonself discrimination. Here, we explore the sequence space of PIWI-interacting RNAs (piRNAs), the genome defense of animals, and establish functional priority among individual sequences for the first time. Our results reveal that only the topmost abundant piRNAs are commonly present in every cell, while rare sequences generate cell-to-cell diversity in flies and mice. We identify a skewed distribution of sequence abundance as a hallmark of piRNA populations and show that quantitative differences of more than a thousand-fold are established by conserved mechanisms of biogenesis. Finally, our genomics analyses and direct reporter assays reveal that abundance determines function in piRNA-guided genome defense. Taken together, we identify an effective sequence space and untangle two classes of piRNAs that differ in complexity and function. The first class represents the topmost abundant sequences and drives silencing of genomic parasites. The second class sparsely covers an enormous sequence space. These rare piRNAs cannot function in every cell, every individual or every generation but create diversity with potential for adaptation in the ongoing arms race with genome invaders.

## INTRODUCTION

Retroviruses and other foreign nucleic acids pose a threat to genome integrity (Kazazian and Moran 2017; Slotkin and Martienssen 2007). In the ongoing arms race with invading nucleic acids, host genomes accumulated scars, eliminated deleterious mutations and selected for the rare advantageous insertions. But above all, they devised defense pathways (Cosby et al. 2019). RNA-guided mechanisms, CRISPR pathways and RNA interference (RNAi), protect the integrity of genomes from bacteria to humans (Koonin 2019; Williams et al. 2015). Animal germ cells employ a specialized RNAi pathway, PIWI proteins and their PIWI-interacting small RNAs (piRNAs), to establish lasting epigenetic restriction of mobile genetic elements (Ozata et al. 2018; Ophinni et al. 2019). Loss of key piRNA pathway genes universally results in sterility of the animal and threatens the survival of the species (Lin and Spradling 1997; Czech et al. 2018; Ozata et al. 2018; Iwasaki et al. 2015).

Specificity of genome defense is imperative, because failing to silence a single parasite or wrongly restricting a single essential host gene is deleterious. Target specificity is determined by complementary base-pairing and places the sequences of piRNAs at the center of self/nonself discrimination (Brennecke et al. 2008; Wasik et al. 2015; Paul 2010; Janssen et al. 2018). Mature piRNAs are ~30 nucleotides in length and generated from hundreds of precursors that can be more than a thousand times their size (Supplemental Fig. S1A). A single precursor can give rise to hundreds or thousands of different piRNAs and is consumed in the process (Wu et al. 2020; Czech et al. 2018; Ozata et al. 2018; Iwasaki et al. 2015). While core mechanisms of piRNA silencing are conserved from flies to mice, the sequences of piRNAs and thus their target repertoire are variable and poorly understood (Zhang et al. 2020; Parhad and Theurkauf 2019; Ozata et al. 2019).

In *Drosophila,* a genomic region of more than 100 kilobases, *flamenco* (flam), has long been known as a major transposon control region (Sarot et al. 2004) (Supplemental Fig. S1B). This essential piRNA cluster looks like a transposon graveyard with densely packed fragments of endogenous retroviruses (Brennecke et al. 2007). It is suggested to produce a single transcript that captures most transposon fragments in antisense orientation, so that the resulting piRNAs identify these very elements by sequence complementarity. Insertion of a novel sequence into a piRNA cluster promotes silencing of complementary targets, and a single fortunate insertion of a genomic invader could provide immunity against the parasite (Yu et al. 2019b; Muerdter et al. 2011; Zhang et al. 2020). Upon association with PIWI-proteins, piRNAs become sequence-specific guides that trigger transcriptional and post-transcriptional restriction (Siomi et al. 2011).

PiRNAs are generated from their long precursors by the conserved endonuclease Zucchini (Zuc)/Pld6 or by the piRNA-guided nuclease activity of PIWI proteins themselves (Ipsaro et al. 2012; Nishimasu et al. 2012; Gunawardane et al. 2007; Brennecke et al. 2007; Czech et al. 2018). The Zuc-processor complex has a characteristic bias to generate piRNAs with a Uridine at the 5’ most position (1U-bias) (Stein et al. 2019; Izumi et al. 2020). Additional sequence motifs, RNA structures, RNA binding proteins, and piRNA-guided cleavage itself have been implicated to instruct patterns of piRNA biogenesis and shape the piRNA sequence space (Pandey et al. 2017; Ishizu et al. 2015; Rogers et al. 2017; Han et al. 2015; Mohn et al. 2015; Izumi et al. 2020). However, a universal signature remains elusive. Here, we identify conserved rules that govern the production of abundant piRNAs, and establish functional priority among individual piRNA sequences for the first time.

## RESULTS

### A single cell contains only a fraction but never the entire set of piRNA sequences

PiRNAs comprise millions of unique sequences in flies and mice (Fig. 1A). The entire piRNA population is even larger than what we can observe in our datasets. Based on species accumulation curves, we predict millions of unique sequences in a saturated population of about half a billion Piwi-piRNAs and hundreds of millions Miwi- or Mili-piRNAs (Fig. 1B) (Deng et al. 2015). The large number of piRNAs presents a biological dilemma since there is a physical constraint on how many piRNAs a cell can accommodate. Using a synthetic reference (Farazi et al. 2011), we estimated that the total number of piRNAs in a single *Drosophila* ovarian somatic sheath cell (OSC) (Saito et al. 2009) does not exceed one million (Fig. 1C and Supplemental Fig. S1C). Our estimates bolster previous observations in mice that place piRNAs among the most abundant molecules in a cell with numbers potentially close to ribosome (Gainetdinov et al. 2018). However, despite the large number of piRNAs within a cell, there are more unique sequences than the total number of molecules that a single cell can contain. Our data imply that each cell contains only a fraction but never the entire set of piRNA sequences. With the essential role of piRNAs in germ cells, the heterogenous complement of single cells could be a key contributor to reproductive polymorphisms and epigenetic variability.

**Figure 1.**
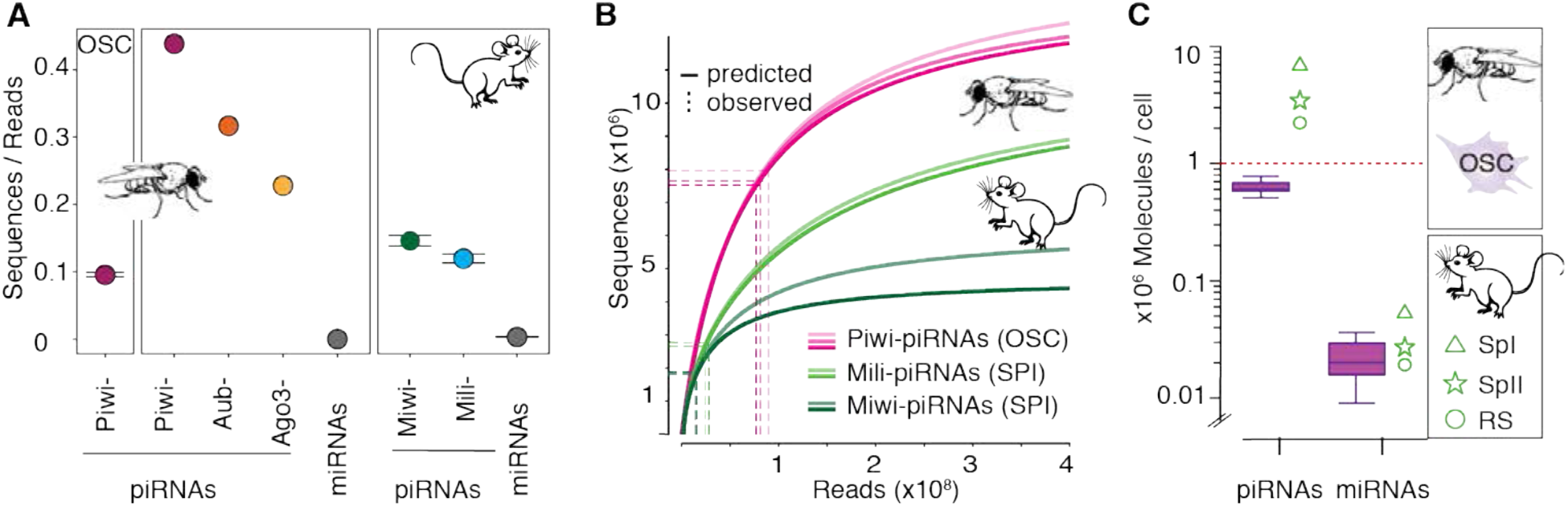
The sequence diversity of piRNAs exceeds the capacity of an individual cell and generates cell-to-cell variability. (**A**) Sequence diversity (Sequences/Reads) of piRNAs associated with different PIWI proteins in flies and mice. Piwi-piRNAs in ovarian somatic sheath cells (OSC) (mean ±SD; n = 3, this study). Piwi-, Aubergine-(Aub) and Argonaute-3-(Ago3) piRNAs in Drosophila ovaries (GEO: GSE83698) (n=1). Miwi/Piwil1- and Mili/Piwil2-piRNAs in primary spermatocytes (SRA: PRJNA421205) (n=2, range indicated) (Supplemental Table S1). For comparison: microRNAs (miRNAs) according to miRbase annotation from total small RNA data sets (GSE83698 and SRR3715418). **(B)** Prediction of piRNA populations according to species accumulation curves based on experimental sampling. Piwi-piRNAs from OSC (this study) (n=3) and Mili- and Miwi-piRNAs from primary spermatocytes (SPI) (n=2) (SRA: PRJNA421205). The number of sampled reads (x-axis) and sequences (y-axis) is indicated by dotted lines. **(C)** The average number of Piwi-piRNAs and miRNAs in a single cell. Numbers based on calibrated sequencing of total small RNAs from OSC (median, 25^th^-75^th^ percentile, n=8) (Fig. S1C). Mouse data from primary spermatocytes (SPI), secondary spermatocytes (SPII) and round spermatids (RS) from (Gainetdinov et al. 2018) are shown for comparison.

### A skewed distribution of sequence abundance results in a few common and many rare piRNAs

Next, we aimed to identify the group of piRNAs that is common to all cells. Based on our estimate that a single ovarian somatic sheath cell (OSC) cannot contain more than one million Piwi-piRNAs, we posit that a piRNA needs to occur more than once in a million to be potentially present in every cell. To track the abundance of individual piRNA sequences, we calculated their concentration (in parts per million, ppm) and ranked them by this measure of sequence abundance (Fig. 2A). Our analysis revealed that the abundance of individual piRNA sequences is highly skewed and varies by more than 1000-fold. Less than five percent of the sequences can be present in every cell (Fig. 2A, red dotted line). Most of the sequences are seen less than once in a million and about half are only represented by a single read in an average dataset (Fig. 2A bottom). The skewed distribution of sequence abundance is a conserved feature of piRNAs in flies and mice based on publicly available data (Gainetdinov et al. 2018; Hayashi et al. 2016), and identifies a small group of abundant sequences that dominate piRNA populations (Fig. 2B and C). In agreement with the small fraction of highly abundant sequences, less than 20% of the topmost abundant Piwi-piRNAs, and about 30% of mouse piRNAs can be commonly found in independent biological data sets (Fig. 2D and E). Our results reveal differences in sequence abundance as a characteristic of piRNA populations, and raises two main questions: Why are some piRNAs so much more abundant than others and how much does abundance matter for function?

**Figure 2.**
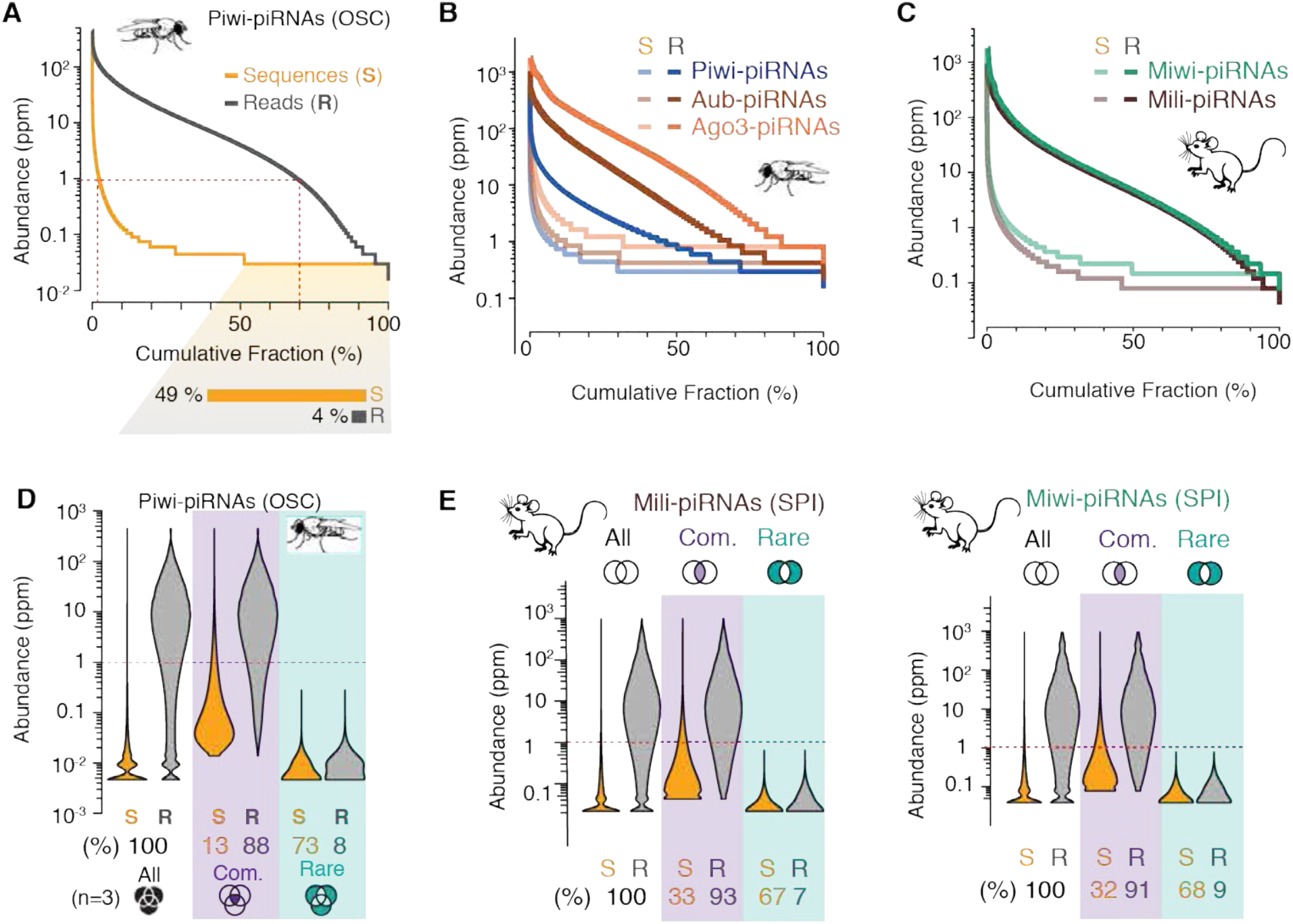
A skewed distribution of sequence abundance results in a few common and many rare piRNAs. The abundance of individual piRNA sequences (Abundance) varies by ~ 1000-fold-ranging from ~0.1 to more than 1000 reads per million (ppm)-in flies (**A, B**) and mice (**C**) (n=1). **(A)** Individual Piwi-piRNA sequences from OSC (S, orange) were ranked by their abundance in reads per million (ppm) (Supplemental Table S2) (n=1). Cumulative distribution of corresponding reads (R, grey). The fraction of sequences that is only represented by a single molecule in a representative data set is indicated below. (**B**) Sequence abundance and cumulative read distribution as in (A) for piRNAs that were associated with Piwi, Aubergine (Aub) and Argonaute-3 (Ago3) in Drosophila ovaries (GEO: GSE83698), and **(C)** Mili/Piwil2 and Miwi/Piwil1 in murine primary spermatocytes (SPI) (SRA: PRJNA421205). **(D)** Only 13% of most abundant sequences can be commonly found in three independent data sets but make up 88% of all sampled piRNAs. Violin plots depict the abundance of individual sequences (S) and cumulative reads (R) for Piwi-piRNAs in three biological datasets. Common piRNA sequences are found in all three replicates (purple). Rare piRNAs are only observed in one of the three samples (teal). Schematic Venn diagrams indicate the intersecting sets (n=3). (Supplemental Table S2). (E) Common and rare piRNAs (analysis as in D) from the intersection of two biological replicates (n=2) for Mili- and Miwi-piRNAs from murine primary spermatocytes (SRA: PRJNA421205)

### Abundant and rare piRNAs originate from the same precursors

With the goal to identify mechanisms that determine the abundance of individual piRNA sequences, we hypothesized that abundant and rare piRNAs either originate from different long precursors or are generated by different processing mechanisms. We observe that about half of all common Piwi-piRNAs originate from piRNA clusters and in particular from the flamenco region (Fig. 3A insert). To our surprise, almost all rare Piwi-piRNAs originate from piRNA clusters too. However, they were not enriched for flamenco-derived sequences. When we systematically compared piRNAs across 450 clusters, we observed that the mean sequence abundance of common piRNAs varies about 100-fold between different clusters and suggest a ranking of these piRNA generating regions (Fig. 3A). Only the top-ranked clusters produced sequences with a mean abundance greater than one in a million and thus the potential to be present in all cells (red dotted line). Our results show that the abundance of individual piRNAs is linked to their genomic origin, and that only a few top-ranked clusters produce highly abundant piRNAs.

**Figure 3.**
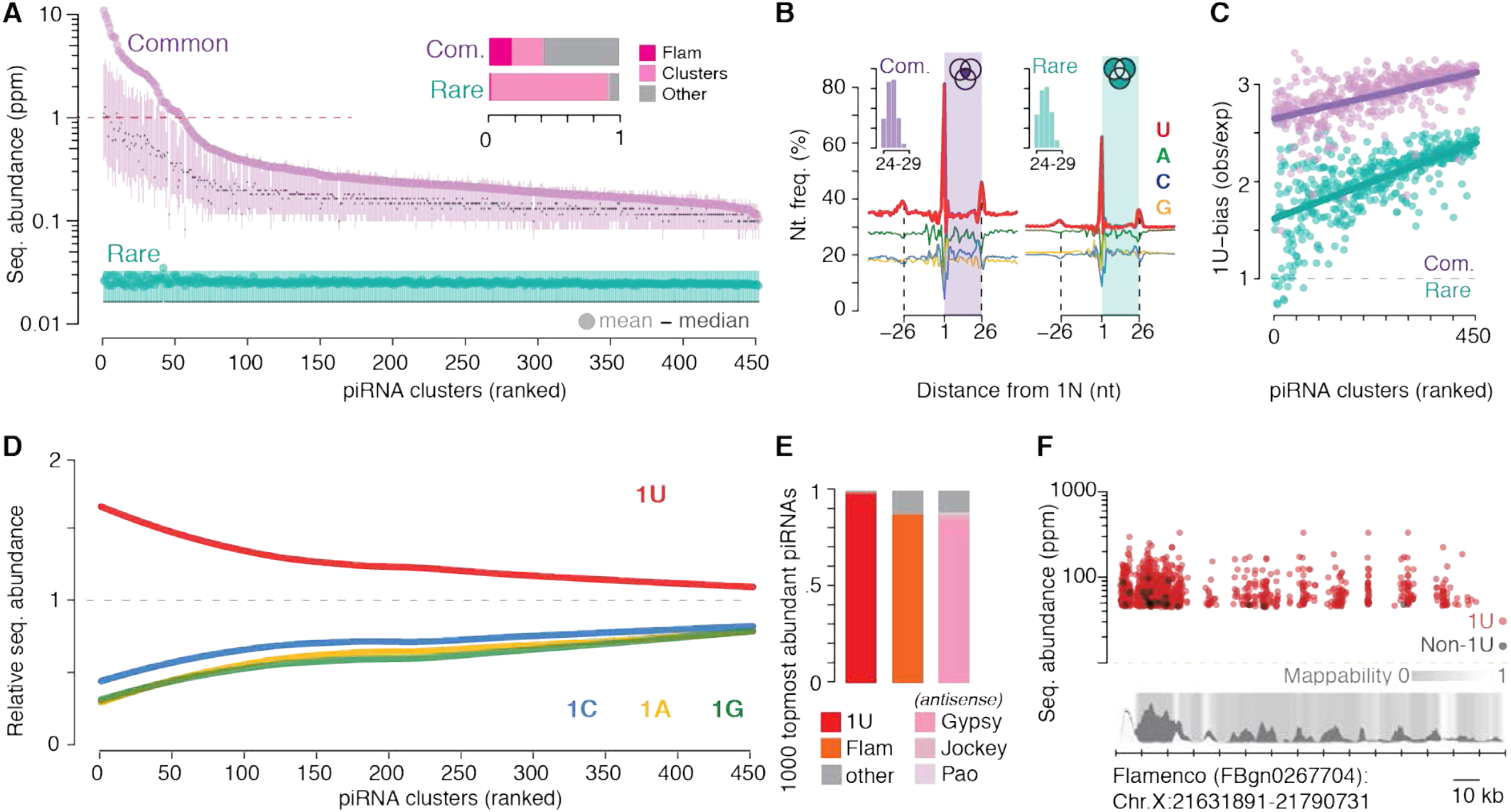
Precursor and processing preferences determine piRNA sequence abundance. **(A)** A small number of top-ranked piRNA clusters (Supplemental table S3), led by flamenco (Flam), produces common sequences (purple) with an average abundance of more than one read per million (ppm) (red dotted line). Clusters were ranked by the mean abundance of common sequences (Supplemental table S3). The sequence abundance of common and rare Piwi-piRNAs for each cluster is shown (mean; median, 25^th^-75^th^ percentile; n=3). Insert: The fraction of common and rare Piwi-piRNAs that originate from Flam and other piRNA clusters. **(B)** Common and rare Piwi-piRNAs are generated by the Zuc-processor. Metagene analyses reveal the phased preference of the Zuc-processor for Uridine (U) in the first position of the observed piRNAs (colored box) as well as in the first position of proceeding and preceding piRNAs. Insert: Length distribution of common and rare piRNAs in nucleotides (nt). **(C)** Common piRNAs have a stronger 1U-bias than rare piRNAs from the same cluster. The bias for Uridine (1U-bias) in the first position-observed over expected 1U frequencies (obs/exp)-of Piwi-piRNAs (clusters ranked as in **A** and **D**). **(D)** Uridine in the first position (1U) is indicative of higher sequence abundance. The mean abundance of piRNAs that start with a Uridine (1U) was compared to that of piRNAs that start with either Adenosine (1A), Guanosine (1G) or Cytidine (1C). The mean abundance of 1U-, 1A-, 1G-, and 1C-sequences relative to the mean abundance of all sequences from the same cluster (relative sequence abundance) is shown (clusters ranked as in **A** and **C**). **E and F,** Most of the 1000 topmost abundant Piwi-piRNAs exhibit a 1U, originate from Flam and show antisense-complementarity to Gypsy endogenous retroviruses. Annotated fractions (**E**) and genomic position within Flam (**F**). Multimapping sequences are represented with all possible coordinates within Flam (**F**).

However, this simple relationship of precursor and product cannot explain the groups of rare piRNAs that originate from all piRNA clusters (Fig. 3A). To test, if abundant and rare piRNAs are produced by different processing mechanisms, we probed for the preference of the Zuc-processor to generate piRNAs with Uridine in the first position (1U) (Stein et al. 2019; Mohn et al. 2015; Han et al. 2015). We observe he characteristic phased 1U-signature for common and rare piRNAs, though the preference for 1U is less pronounced in the rare group (Fig. 3B). When we normalized the observed to the expected 1U frequencies (1U-bias) for each cluster, we observed that rare piRNAs consistently exhibit less bias for Uridine in the first position than abundant piRNAs from the same piRNA precursor (Fig. 3C). The differences in the 1U-bias and in sequence abundance were particularly pronounced for piRNAs that originate from the top-ranked piRNA clusters.

### Sequence preferences modulate abundance

Indeed, when we grouped piRNA sequences solely by the identity of their first nucleotide, we observed that the presence of a 1U alone was indicative of higher sequence abundance with the biggest differences for the top-ranked piRNA clusters (Fig. 3D). Among Non-1U-piRNAs, sequences with Adenosine (A), Cytidine (C) or Guanosine (G) in the first position were equally lower in abundance. Taken together, our results suggest that the topmost abundant piRNAs originate from a few top-ranked precursors and harbor a Uridine in the first position. To test this hypothesis, we characterized the top 1000 most abundant Piwi-piRNAs (Fig. 3E, F). 98% of these top 1000 sequences start with Uridine and 88% can be generated by a single piRNA cluster, *flamenco,* the only known essential piRNA cluster (Sarot et al. 2004). More than 80% of these 1000 topmost abundant sequences show antisense complementarity to endogenous retroviruses of the gypsy family in accordance with the known function of *flamenco* in controlling these elements (Sarot et al. 2004; Brennecke et al. 2007). Our results reveal that the abundance of individual piRNAs is determined by their genomic origin and by the identity of their first nucleotide and suggest that the abundance of individual piRNAs is regulated during piRNA biogenesis.

### A conserved mechanism discriminates abundant from rare piRNAs in flies and mice

We observe similar signatures for mouse Mili- and Miwi-piRNAs from primary spermatocytes mining publicly available data from the Zamore group (Gainetdinov et al. 2018) (Fig. 4A). Both, common and rare piRNAs originate from the same pachytene piRNA precursors but exhibit marked differences in mean sequence abundance. Both piRNA groups exhibit a 1U signature, with reduced 1U-preference in rare piRNAs. Indeed, the 1U-bias of rare piRNAs is generally lower than that of common piRNAs (Fig. 4B). Both groups, abundant and rare piRNAs, exhibit the characteristic 3’-end processing signatures of murine piRNAs including Pnldc1-dependent trimming and the +1U-signature of the pre-processing event, likely generated by the murine (m)Zuc-processor (Gainetdinov et al. 2018) (Supplemental Fig. S2A and B). These results suggest that abundant and rare piRNAs are generated by the same processing mechanisms.

**Figure 4.**
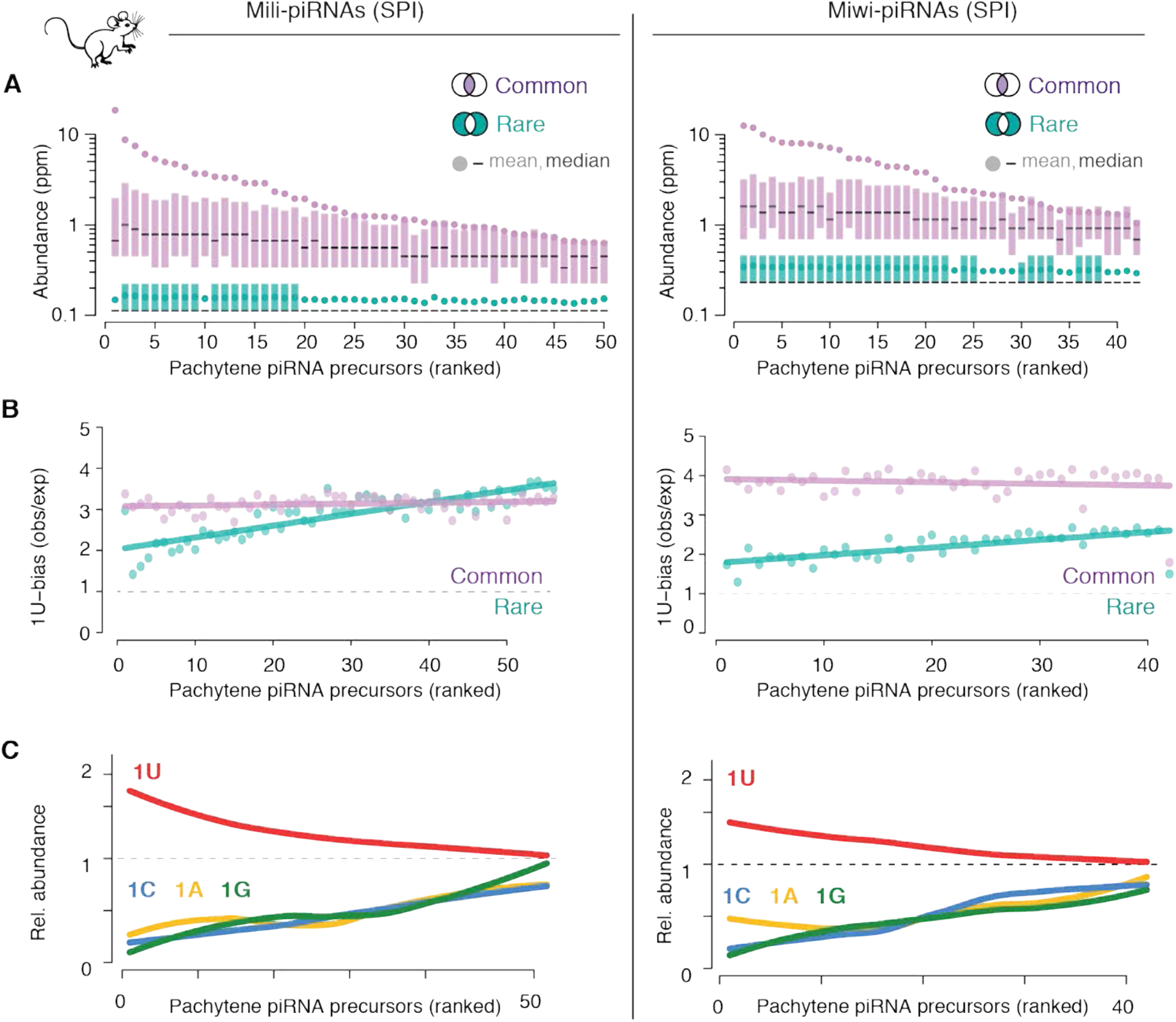
Precursor and processing preferences determine piRNA sequence abundance in mice. **(A)** PiRNA precursors can be ranked by the mean sequence abundance of common piRNAs. All precursors, even top-ranked ones also produced rare piRNAs. The sequence abundance of common and rare piRNAs associated with Mili or Miwi in primary spermatocytes (SPI) is shown for individual pachytene piRNA precursors (mean; median, 25^th^-75^th^ percentile; n=2). Publicly available source data: SRA: PRJNA421205. **(B)** Common piRNAs have a stronger 1U-bias than rare piRNAs from the same precursor. The bias for Uridine (1U-bias) in the first position-observed over expected 1U frequencies (obs/exp)-of Miwi- and Mili-piRNAs is shown for common and rare sequences from individual pachytene piRNA precursors (precursors are ranked as in **A**). **(C)** Uridine in the first position (1U) is indicative of higher sequence abundance. The mean abundance of piRNAs that start with a Uridine (1U) was compared to that of piRNAs that start with either Adenosine (1A), Guanosine (1G) or Cytidine (1C). The mean abundance of 1U-, 1A-, 1G-, and 1C-sequences relative to the mean abundance of all sequences from the same cluster (relative sequence abundance) is shown (pachytene piRNA precursors are ranked as in **A** and **B**).

Finally, we asked, if the 1U-bias determines differences in piRNA sequence abundance also in mice. We grouped all piRNAs from each precursor by their first nucleotide into 1U-, 1C-, 1A- and 1G-piRNAs and calculated their relative abundance. Like for *Drosophila* Piwi-piRNAs, the presence of a 1U alone correlates with increased sequence abundance for Mili- and Miwi-piRNAs. Overall, our comprehensive cross-species analyses identified a conserved signature that discriminates abundant from rare piRNAs. Our data show that piRNA abundance depends on processing preferences and suggest that the position of Uridines across precursors shapes the composition of mature piRNA populations. This simple conserved mechanism could enable upregulation of essential piRNAs and suppression of auto-aggressive sequences during purifying selection.

### Abundance determines function

While it is obvious that piRNAs that cannot be present in every cell cannot act in every cell, we wanted to know, if more subtle changed in piRNA abundance affect piRNA function. To directly measure how much piRNA abundance impacts piRNA-guided silencing, we developed a reporter assay. Our design aimed at providing a quantitative readout for piRNA-mediated silencing at single cell level. We placed target sites with complementarity to endogenous piRNA-generating regions in the 3’ UTR of a green fluorescent protein (GFP) and expressed a red fluorophore (mCherry) from the same plasmid as normalization control (Fig. 5A). Both fluorophores were driven by the same minimal promoter and separated by an insulator element to avoid spreading of silencing. We transfected our sensor into ovarian somatic sheath cells (OSC) and evaluated the expression of both fluorophores after 48 hours. In the absence of piRNA-targeting, both fluorophores were expressed, and most cells appeared yellow (Fig. 5A). We anticipated that repression of GFP by endogenous piRNAs would result in ‘red-only’, GFP-silenced, cells. To quantify the silencing effect at single cell resolution, we analyzed cells by flow cytometry. In the absence of a piRNA-target-sequence, 76% of the cells expressed both green and red fluorophores (Fig. 5B). As expected, a 460nt long target with antisense (as) complementarity to *flamenco* (Flam(as)-460) resulted in decreased green fluorescence and 71% of the cells appeared fully silenced (‘red-only’) (Fig. 4C, and Supplemental Fig. S4A). When this Flam target-sequence was split in half (Flam(as)-230-A and B), either part resulted in about 50% fully silenced cells (Supplemental Fig. S3A-C). Shortening the target region to 100nt (Flam(as)-100), further reduced the occurrence of red-only cells to 2% (Supplemental Fig. S3D). Sensors with 230nt complementarity to the piRNA-producing regions of *cg17514, Traffic jam* (Tj), *Pathetic* (Path) and *Cyclin B* (CycB) resulted in up to 33% fully silenced cells (Supplemental Fig. S3E-I). Overall, we observed a range of piRNA-guided restriction for different sensors from a barely measurable effect to almost complete silencing.

**Figure 5.**
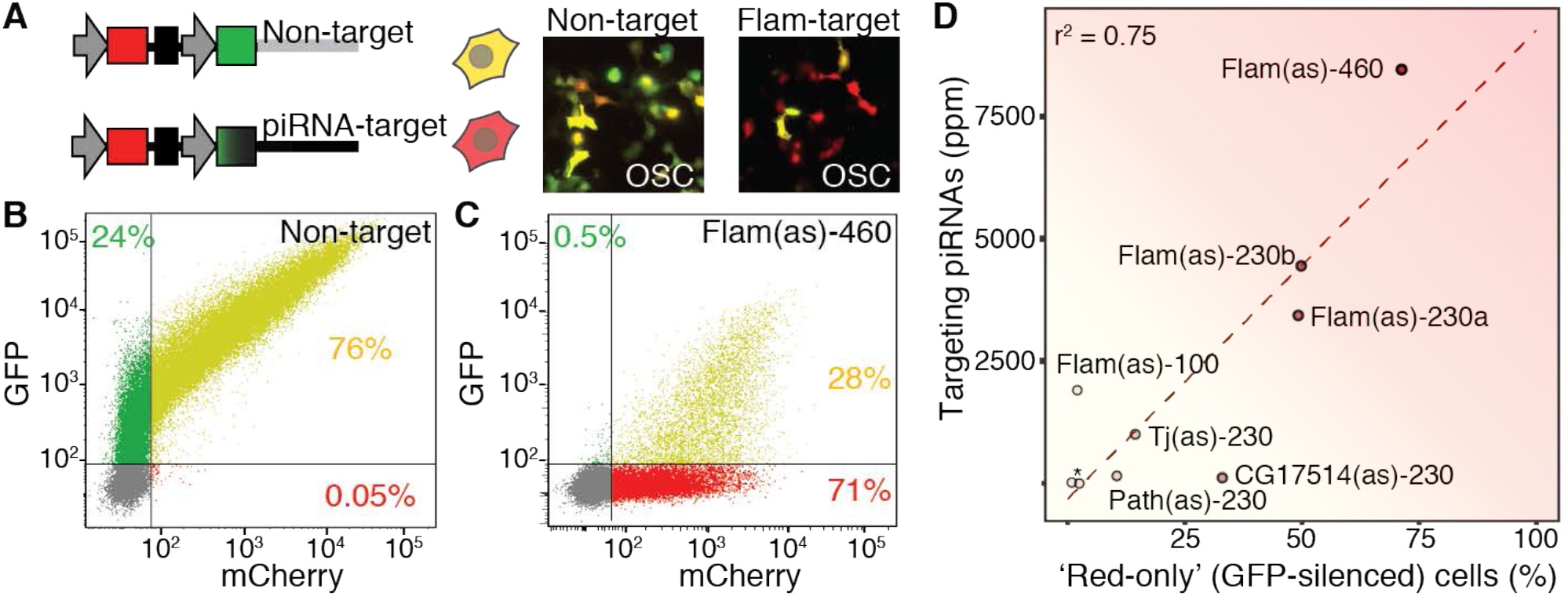
Silencing directly correlates with piRNA abundance. **(A)** A reporter assay for Piwi-piRNA silencing. GFP (green) reports silencing by endogenous piRNAs. mCherry (red) serves as control. GFP and mCherry are expressed from the same plasmid by individual promoters and separated by an insulator sequence. Different target sites with antisense complementarity to piRNA generating regions were inserted into the 3’UTR of GFP (piRNA target) (Supplemental Table S4). Sensors were expressed in ovarian somatic sheath cells (OSC). Expression of the dual color reporter was visualized (A) and measured by flow cytometry (B, C) 48h after transfection. (**B**) Without any target site (non-target), the sensor expressed both fluorophores in 76% of the transfected cells. (**C)** A Flam-target sensor (Flam(as)-460) showed complete silencing of GFP in 71% of the transfected cells (‘red only’ cells). **(D)** PiRNA abundance correlates with silencing. Correlation of ‘red-only’ (GFP-silenced) cells and the total abundance of complementary piRNAs (in parts per million, ppm). Pearson’s Correlation Coefficient (r^2^). PiRNA-sensors with different target sequences complementary to Flam (varying target length: 100, 230, 460nt) or complementarity to other piRNA-producing regions: Traffic jam (Tj), Pathetic (Path), CG17514, and two different intervals of Cyclin B (*) (constant length 230nt) (Supplemental Table S4).

We next asked, whether the abundance of complementary endogenous piRNAs influenced the fraction of fully silenced, ‘red only’, cells (Fig. 5D). We observed a positive correlation between the total number of complementary piRNAs and the number of fully silenced cells (Pearson correlation coefficient, r^2^=0.72). Our results suggest that the combined abundance of complementary piRNAs determines the efficacy of target restriction.

## DISCUSSION

Overall, a model emerges that attributes the significance of individual piRNA sequences to their abundance, and abundance to mechanisms of piRNA biogenesis (Fig. 6). The topmost abundant sequences originate from a few piRNA-generating regions and are characterized by preferences of the Zuc-processor, resulting in a strong 1U-bias. This class of piRNAs represents only a small fraction of the observed sequence space but dominate the functional piRNA molecules in every cell. This outstanding group of *Silencers* has the potential to define nonself for future_generations. An intermediate group of *Modifiers* contains sequences that are not present in every cell but are commonly found in different animals. These piRNAs could collaborate and modulate the efficacy of piRNA-guided restriction on converging targets. These *Modifiers* establish functional cell-to-cell diversity and could contribute to previously observed polymorphisms in piRNA-guided restriction (Ryazansky et al. 2017). Finally, innumerous piRNA sequences are extremely rare. They seem to be functionally insignificant at first glance. However, these *Junk* piRNAs could provide substrate for evolutionary tinkering (Palazzo and Koonin 2020). The sequences themselves or the mechanism that generates them present an opportunity for purifying selection to revise the arsenal of piRNA ammunition in response to a novel genome invader (Yu et al. 2019a).

**Fig. 6.**
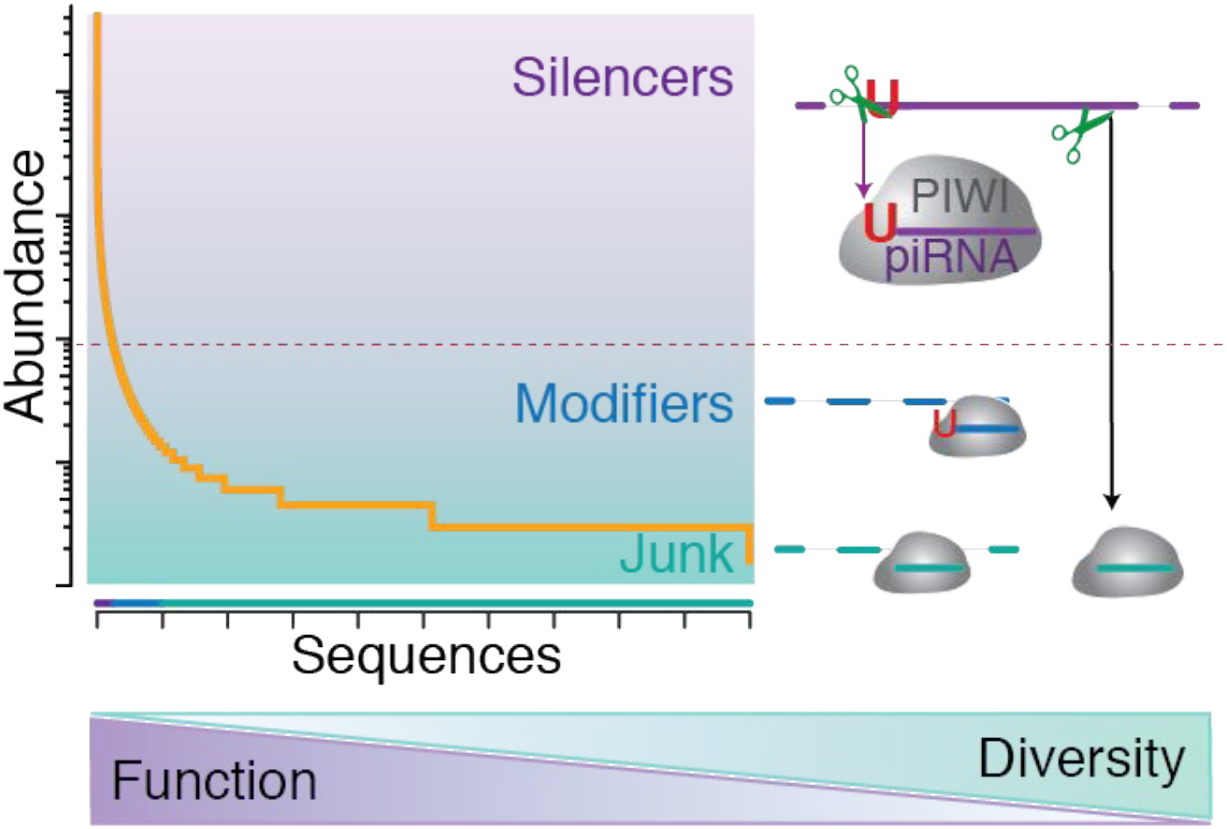
Two classes of piRNAs separate silence from diversity. . Based on the limited number of piRNAs in a single cell (Fig. 1) and the skewed distribution of their sequence abundance (Fig. 2), only the topmost abundant piRNA sequences can be present in every cell. These commonly detected sequences originate from a few top-ranked piRNA generating regions and exhibit a strong preference for Uridine at their 5’ most position (1U) (Fig. 3 and 4). The 1U-preference of the Zuc-processor complex modulates the abundance of individual piRNAs (Fig. 3). Abundance correlates with function (Fig. 3E, F and 5). Based on their sequence abundance, piRNAs can be divided into functional classes. A few topmost abundant sequences dominate piRNA silencing *(Silencers).* A biological threshold (red dotted line) separates these piRNAs from the bulk of low abundant sequences that cannot be present in every cell. Rare piRNAs establishes cell-to-cell diversity. On convergent targets, these piRNAs could act as *Modifiers* and promote reproductive polymorphism. In isolation, extremely low abundant piRNAs might never act. However, over time, these highly diverse *Junk* sequences could serve as resource for evolutionary tinkering and bolster adaptation to novel genomic invaders. The functional sequence space of piRNAs is concise and can be regulated to ensure careful self/nonself discrimination.

Self/nonself discrimination is at the heart of every self-defense, and regulatory mechanism are required to adjust the dial. In the ongoing arms race with genome invaders, the functional piRNA sequence space requires adjustments to restrict nonself and avoid auto-aggression. Function requires abundance, and only the most abundant piRNAs efficiently restrict complementary targets. Regulated abundance ensures a concise functional sequence space. Throughout evolution, purifying selection could act on entire piRNA generating regions and within individual precursors to shape the repertoire of functional piRNAs for maximized genome protection and minimized auto-aggression. The plethora of rare *junk* sequences could serve as substrate for evolutionary tinkering and successful adaptation to novel genomic residents.

## METHODS

### Purification of Piwi-piRNAs

Piwi-piRNAs were extracted from ovarian somatic sheath cells (OSC) by immunoprecipitation of endogenous or endogenously Flag-tagged Piwi (eF-Piwi): The anti-Piwi antibody and Surebeads Protein A Magnetic bead (Bio-Rad, 1614013) were used to immunoprecipitated endogenous Piwi-piRNAs from witld type OSC. Anti-Flag M2 magnetic beads (Sigma-Aldrich, M8823) were used to immunopurify eF-Piwi from OSC_eF-Piwi_ (https://doi.org/10.1101/2020.12.18.423517). Three biological replicates were prepared for each sample type. Cells and plasmids are available through the Drosophila Genomics Resource Center (DGRC) and upon request.

### Calibrated total small RNA (tsRNA) sample preparation for Illumina sequencing

Calibrated small RNAs were prepared for the quantification of small RNAs (eight biological replicates, n=8). The samples were prepared following procedures described in Hafner et.al. 2012 (PMID: 22885844) with the following modifications: First, total small RNAs from one million OSC cells were extracted using PureLink miRNA Isolation kit (Thermo Fisher Scientific, K157001) and were spiked with a calibrator mix (see below for preparation, calibrators 1-4, Supplemental Table S5) (Supplemental Fig. 1C). The calibrated small RNA samples were ligated to an adenylated and fluorescently labeled 3’ adaptor using T4 RNA ligase 2 truncated KQ (NEB, M0373S) and were subsequently separated on a 12% polyacrylamide UREA gel. Adaptor ligated RNAs 48-58nt long (corresponding to 19-29nt long input RNAs) were extracted and ligated to the 5’ adaptor using T4 RNA Ligase 1 (NEB, M0204). A total of ten variable nucleotides (unique molecular identifiers, UMIs) were included in the adaptor sequences (eight in the 5’ adaptor and two in the 3’ adaptor) (Supplemental Table S5). The 10 UMIs were used for removal of PCR duplicates during analysis. After 5’ ligation, RNA was recovered using Oligo Clean and Concentrator kit (Zymo Research, D4060) and used as template for cDNA synthesis using SuperScript IV First-Strand Synthesis System kit (Invitrogen, 18091050). The generated cDNA was amplified by PCR and the libraries were purified on a Pippin Gel Cassette (3% Agarose w/ Ethidium Bromide 40 μl EM, Sage Science) using the Pippin Prep system (Sage Science). Quality of the samples was assessed on a DNA Screen Tape (Agilent, D1000) using a TapeStation (Agilent). The calibrated small RNA samples were sequenced using an Illumina NextSeq 550 and 50nt long single end reads were obtained.

### Dual color reporter assay: Plasmid design and construction

To create the dual-color reporter plasmid, the sequence of mCherry-T2A-(Puromycin-resistance) was retrieved from pCDH-CMV-mCherry-T2A-Puro (Addgene plasmid # 72264) and was cloned into pUC19 at the HindIII cleavage site using HiFi assembly (NEB, E2621S). The physical DNA sequence of gypsy insulator was a gift from Dr. Brian Oliver’s group (NIDDK) (sequence as in pCFD6; 73915, Addgene) and was inserted downstream of the mCherry-T2A-Puromycin resistance stretch using restriction cloning (AscI-XhoI). EGFP was synthesized as a g-block by Integrated DNA Technologies (IDT) and was inserted downstream of the gypsy insulator using HiFi assembly at the XhoI cleavage site. Both the EGFP and the mCherry are driven by a 595-nt sequence upstream of Piwi gene that contains the core Piwi promoter. The sensor sequences corresponding to complementary sequences of piRNA-generating regions were synthesized as g-blocks by IDT and ligated into the 3’ UTR of EGFP at the SrfI restriction site (Fig. 5, Supplemental Fig. S3 and Supplemental Table S4).

### Data analysis

was carried out mostly in the R (v3.6.2) statistical framework using following Bioconductor packages: Rsamtools (2.2.3), BSgenome (1.54.0), GenomicRanges (1.38.0), GenomicAlignemnts (1.22.1), Biostrings (2.54.0), data.table (1.12.8), ggplot2 (3.3.1), stringr (1.4.0), reshape (0.8.8), reshape2 (1.4.4), plyr (1.8.6), dplyr (1.0.0), rlist (0.4.6.1), Rsubread (2.0.1), ShortRead (1.44.3), Gviz (1.30.3), ggseqlogo (0.1).

**Detailed molecular and computational methods are described in the Supplemental material.**

## Data access

All raw and processed sequencing data generated in this study have been submitted to the NCBI Gene Expression Omnibus (GEO; https://www.ncbi.nlm.nih.gov/geo/) under accession number GSE156058. Key count tables are provided as supplemental tables (S1-4). Computational methods are described in methods and custom R scripts are available in on GitHub.

## Acknowledgements

We thank VG Cheung, S Gottesman, and TS Macfarlan for critical comments on the manuscript; M Hafner, NR Guydosh and all members of the Haase, Hafner and Guydosh labs for discussions, especially D Anastasakis and K Bettridge for experimental help, and S Mitra for technical assistance with initial data analyses; the NHLBI flow cytometry and genomics cores, especially A Saxena, the NIH high-performance computing group, and NIH Medical Arts, especially E He for help with the model figure. This work was supported by the intramural research program of the NIDDK.

## Author contributions

P.G. and P.K. and D.S. performed experiments and analyzed data with help from A.M., C.M.A., A.R.E., and C.S.; A.D.H. conceived the project and wrote the manuscript with help from P.G., P.K. and D.S.

